# Changes in social cohesion in a long-lived species under a perturbation regime

**DOI:** 10.1101/686451

**Authors:** M. Genovart, O. Gimenez, A. Bertolero, R. Choquet, D. Oro, R. Pradel

## Abstract

1. Understanding the behaviour of a population under perturbations is crucial and can help to mitigate the effects of global change. Sociality can influence the dynamics of behavioural processes and plays an important role on populations’ resilience. However little is known about the effects of perturbations on the social cohesion of group-living animals.
2. To explore the strength of social cohesion and its dynamics under perturbations, we studied an ecological system involving a colonial, long-lived species living in a site experiencing a shift to a perturbed regime. This regime, caused by the invasion of predators, led this colony to hold from 70% to only 3% of the total world population in only one decade. Because birds breed aggregated in discrete and annually changing patches within large colonies, we could disentangle whether annual aggregation was random or resulted from social bonding among individuals. Our goals were 1) to uncover if there was any long-term social bonding between individuals and 2) to examine whether the perturbation regime affected social cohesion.
3. We explored social cohesion by means of contingency tables and, within the Social Network Analysis framework, by modeling interdependencies among observations using additive and multiplicative effects (AME) and accounted for missing data. We analysed 25 years of monitoring with an individual capture-recapture database of more than 3,500 individuals.
4. We showed that social bonding occurs over years in this species. We additionally show that social bonding strongly decreased after the perturbation regime. We propose that sociality and individual behavioural heterogeneity have been playing a major role driving dispersal and thus population dynamics over the study period.
5. Perturbations may lead not only to changes in individuals’ behaviours and fitness but also to a change in populations’ social cohesion. The demographic consequences of the breaking down of social bonds are still not well understood, but they can be critical for population dynamics of social species. Further studies considering individual heterogeneity, sociality and different types of perturbations should be carried out to improve our understanding on the resilience of social species.

## Introduction

Ecosystems are subject to perturbations, both natural and human induced, affecting individuals, populations and communities. When they are strong or are maintained through time, these perturbations may cause a shift in individual or population states or phases and even lead to collapses and extinctions (Dai, Korolev, & Gore, 2015; Dakos, Carpenter, van Nes, & Scheffer, 2014). Understanding how individuals and populations will respond to these perturbations is critical both from a ‘pure’ ecological standpoint and also from an applied point of view to mitigate the effects of global change (Colchero et al., 2018; Coulson et al., 2017; Donohue et al., 2016). Population dynamics may be directly affected by these perturbations through a decrease in demographic parameters such as survival or fecundity, or by a change, immediate or delayed, on individual behaviour, such as an increase in dispersal (Fernandez-Chacon et al., 2013). We define population resilience as the maximal pulse perturbation a population can tolerate or absorb without going extinct (Dai et al., 2015; Holling, 1973). In social animals, social behavioural processes, such as information sharing and decision-making, add another dimension to understanding the resilience of populations facing perturbations. For instance, the amount of social information can be enhanced not only by positive density-dependence, but also by social cohesion (Barrett, Henzi, & Lusseau, 2012; Centola, 2018; Kerth, Perony, & Schweitzer, 2011). Social bonds favor the exchange of private information and consequently reduce uncertainty in resource acquisition (e.g. shelter against predators, food, mates) or in decision-making in the face of disturbances, such as dispersal to non-perturbed or less perturbed sites (Fernandez-Chacon et al., 2013; Snijders, Blumstein, Stanley, & Franks, 2017; Webber & Vander Wal, 2018). Thus, the structure of a group affects social interaction, information transfer, and collective decisions (Sueur & Mery, 2017). However, little is known about the effects of perturbations on the social cohesion of populations in empirical studies of social animals.

The analysis of social relationships in animal populations may include from simple and ephemeral contacts, to permanent and strong bonds between individuals (Firth & Sheldon, 2015; Genton et al., 2015; Kerth et al., 2011; Leu, Farine, Wey, Sih, & Bull, 2016). Coloniality is a life-history strategy where individuals show a clear social relationship among conspecifics, breeding in large and dense groups (Brown, 2016; Rolland, Danchin, & de Fraipont, 1998). However, many colonial species are philopatric, thus this relationship may not necessarily reflect individual social bonds but a shared tendency to breed in the same birthplace. This tendency may result from the need to share information about resources, especially when they are patchy and more unpredictable, or it may result from the advantages of social defence against predators (Clode, 1993; Hoogland & Sherman, 1976). A challenge lies in disentangling whether annual association between individuals is only due to philopatry, or also due to the existence of a social bonding within groups of individuals over time. If the latter is true, then this would suggest the evolution of social cohesion for exploiting the evolutionary advantages of social living (including social information sharing) for individual fitness prospects.

Social network theory, originated in sociology and widely used to study human relationships and social organization (Centola, 2018; Moreno, 1934; Scott, 1988) now provides both a conceptual framework and the analytical tools to explore social cohesion and social processes in animal populations (Croft, James, & Krause, 2008; Farine & Whitehead, 2015; Ward & Webster, 2016; Whitehead, 2014). Network theory is now being simultaneously developed in a number of fields, including statistical physics, sociology, molecular biology, and computer science. As a result the field is changing at a rapid pace. While not all developments can or should be applied toward the study of animal societies (James, Croft, & Krause, 2009), this rush of novel ideas from outside disciplines is enriching behavioural ecology (Hasenjager & Dugatkin, 2015).

To explore the existence of social cohesion and its dynamics under perturbations, we studied an ecological system involving a colonial, social vertebrate (the Audouin’s gull *Larus audouinii*) living in a site experiencing a shift to a perturbed regime. Interestingly from a social point of view, the species breeds aggregated in spatially-discrete patches within large colonies. Each breeding season, some patches go extinct and some are colonized, forcing individuals to breed in patches different from the ones they were born in or they bred in the previous year. These colonization-extinction processes may allow us to disentangle whether social aggregation among individuals is an annual random association, or it rather results from social cohesion among individuals. An extensive long-term monitoring program has been carried out since 1988 at the Ebro Delta, comprising the main breeding site for the species, the Punta de la Banya (M. Genovart, Oro, & Tenan, 2018). Here, population dynamics has undergone different phases: an initial growing phase after site colonization, an exponential growth phase, a stable phase of dynamic equilibrium, and a final transition phase to population collapse (Almaraz & Oro, 2011; Figure 1). This collapse was due to the arrival of terrestrial predators, and led this colony to hold from 70% to only 3% of the total world population in only a decade (Figure 1; M. Genovart et al., 2018; Payo-Payo et al., 2018). The perturbation regime caused changes in the spatial distribution of patches at the site, changes in age structure, decrease in fecundity and an increase of dispersal to other sites (Payo-Payo et al., 2017, 2018).

**Figure 1.**
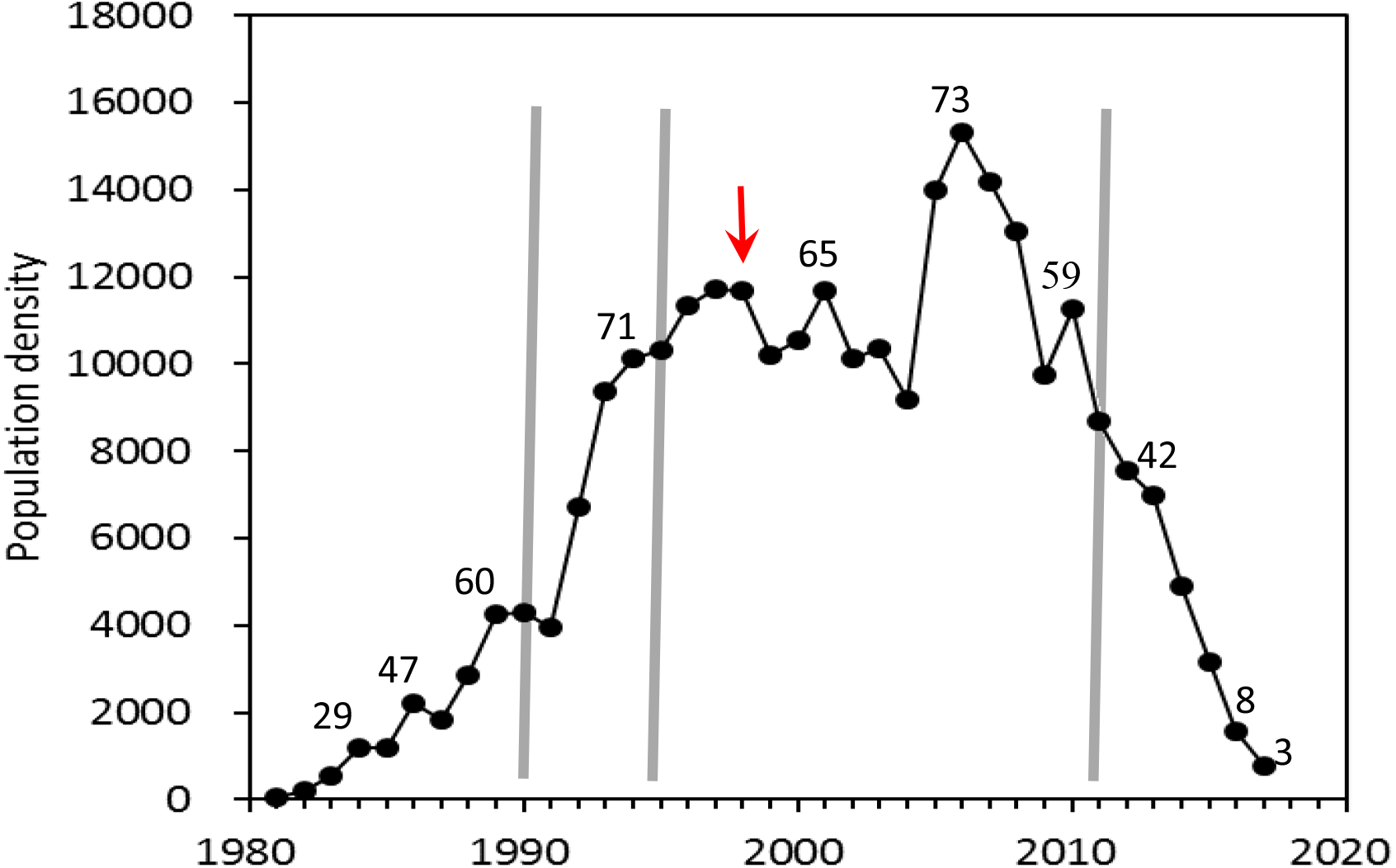
Number of breeding pairs in the Punta de la Banya colony from colonization in 1981 to 2017. The observed phases in the population dynamics (initial growth, dynamic equilibrium and collapse) are separated by grey lines, which are identified by chronological clustering analysis (Almaraz & Oro, 2011). Red arrow indicates the arrival of predators to the colony; a resistance phase started with this invasion and ended with the onset of the transition phase to collapse. Numbers above some years show the percentage of total world population held at Punta de la Banya.

Taking advantage of the long-term monitoring of this long-lived species, the knowledge of its population dynamics, and the use of tools recently developed in the Social Network Analysis (SNA) framework, we specifically addressed the following questions: 1) is there any long-term social bonding between individuals breeding in the same patch and 2) have perturbations, in this case a perturbation regime, affected social cohesion? We finally discuss the role and consequences of social cohesion in population dynamics and resilience in social species.

## Material and Methods

### Study species and study area

The Audouin’s gull is a long-lived seabird with more than 80% of the global population breeding in the western Mediterranean (http://www.iucnredlist.org/details/22694313/0; M. Genovart et al., 2018). This species was critically endangered until the early 80’s, when it colonized a new site, the Punta de la Banya in the Ebro Delta (Figure 2). Here, the large availability of both suitable breeding habitat and food resulted in a rapid and exponential growth, ending with the site holding more than 70% of the total world population in 2006. The global population dynamics was mainly driven by this colony and after the exponential growth, the species was downgraded to a conservation category of “least concern” (IUCN 2015). However, the Punta de la Banya colony is now collapsing and even if the species is colonizing new sites, the global population is decreasing at a 5% annual rate (M. Genovart, Oro, & Tenan, 2018; Figure 1). In 1997, first carnivores (mainly foxes, but also badgers, beech martens and least weasels) arrived at Punta de la Banya, and since then the site has been perturbed to a greater or lesser extent by carnivores.

**Figure 2.**
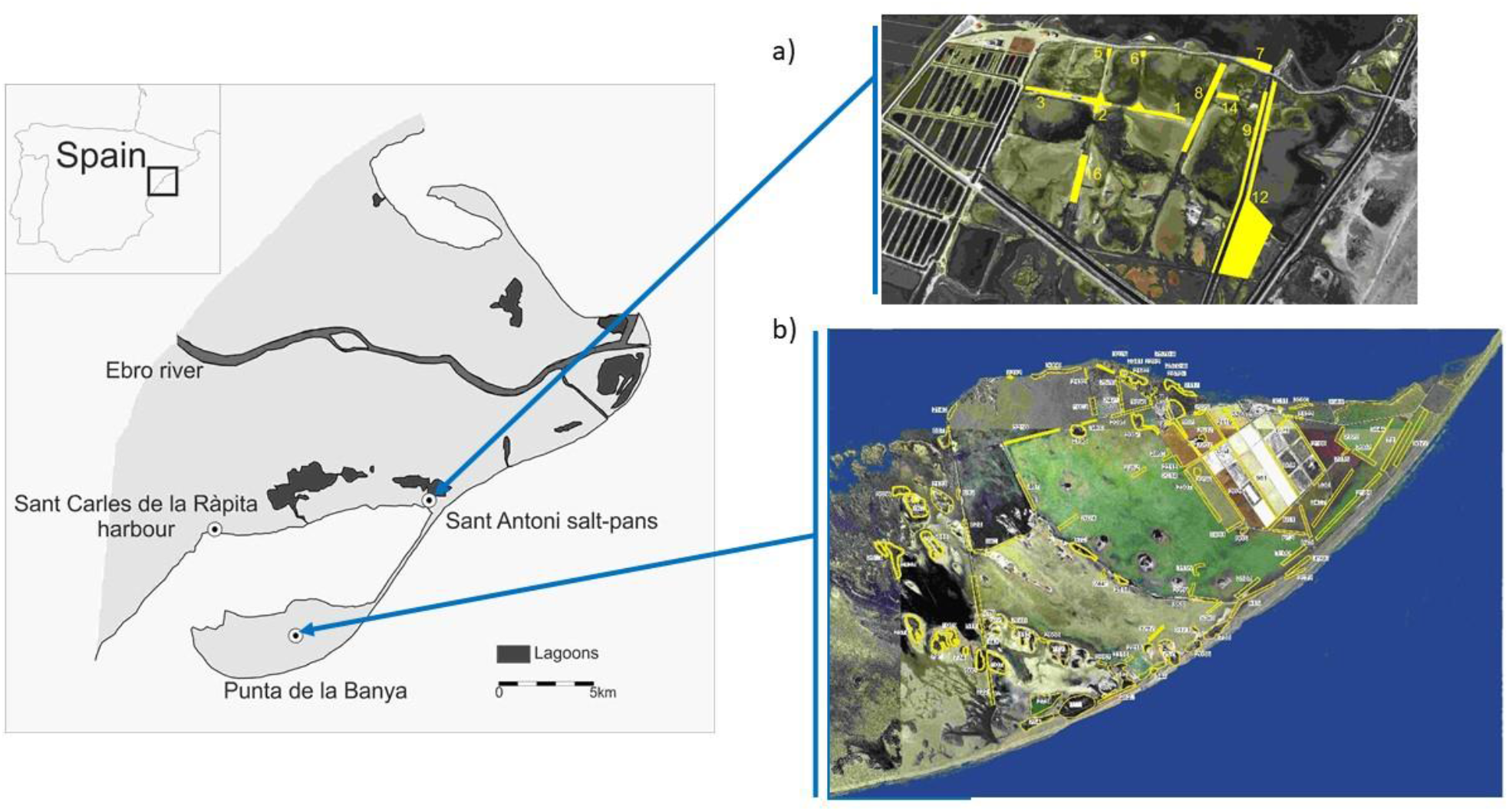
Map of the study area comprising the 3 main colonies and the distribution of patches within colonies during the study period at a) Sant Antoni and b) Punta de la Banya. Sant Carles de la Rápita colony is considered to have only one patch.

Annual censuses of breeding pairs at every patch within colonies at the Ebro Delta area have been carried out since colonization in 1981 to 2017 (Figure 1 and 2; Figure S2). In the Ebro Delta there are three colonies: Punta de la Banya, colonized in 1981 and occupied throughout the study period, Sant Carles de la Ràpita harbour, occupied since 2011 to 2015, and Sant Antoni, occupied from 2013 up to now (Figure 2, Figure S1). Within colonies, individuals are distributed in patches or sub-colonies (Meritxell Genovart, Jover, Ruiz, & Oro, 2003, Figure 2). As patch location may change from one year to another, we annually geolocalized, mapped and defined the breeding patches (Figure 2, Figure S1).

### Individual data

During 1988-2017 a total of 30,290 individuals were captured and ringed as chicks at the Punta de la Banya (Meritxell Genovart, Pradel, & Oro, 2012; Oro, Tavecchia, & Genovart, 2010). From 2002 to 2017 resightings were made using spotting scopes from a distance all over the western Mediterranean with a total of 63,106 resights in the study area and 5,593 different individuals resighted. Each year we recorded the breeding patch for each individual. To make sure that individuals were breeding and that they did so in a particular patch, we only selected those individuals seen during the breeding season in a particular patch showing unequivocal breeding behaviour. Specifically, individuals making alarm calls, incubating eggs or with chicks. After this selective filter our final database included 3,548 individuals.

### SNA framework

Our social network, defined as the observed pattern of breeding association, was constructed taking individuals as the nodes of the network and each edge dyad (i.e. pair of individuals) representing the fact that individuals breed in the same patch. We ended with a global sociomatrix, i.e. the matrix representation of the dyadic relationships among individuals, of 3,548* 3,548. Depending on the question addressed, edges showed if two individuals bred in the same patch one year, or at least once in a certain period (see below). The network was not directional. The strength of association between dyads was calculated using the half-weight association index (HWI) more suitable when not all individuals within each group have been identified (Cairns & Schwager, 1987; Ginsberg & Young, 1992). In our case, the HWI measures the proportion of time individuals have bred together, from 0 (never bred together) to 1 (never observed breeding apart). Based on previous results on population dynamics, and on the population size of this species and colony (M. Genovart et al., 2018; Payo-Payo et al., 2017), we divided our dataset in two main periods: a period defined as “stable phase” from 2002 to 2011, and a period of “transition phase to collapse”, from 2012 to 2017 (Figure 1).

For some of our analyses we used the recently developed AME function from the AMEN package (Hoff, 2018; Minhas, Hoff, & Ward, 2016) that can be applied to binary, ordinal, and continuous network data. This new approach uses an iterative Markov chain Monte Carlo (MCMC) algorithm that provides Bayesian inference of the parameters in the social relations regression model (SRM; Warner, Kenny, & Stoto, 1979) using additive and multiplicative effects and combining the linear regression model with the covariance structure of the SRM (Minhas et al., 2016). The AME method is also able to cope with missing and censored data; this is highly relevant when analysing sociality on wild populations, as detection rate for individuals is almost always imperfect, and properly controlling for missed observations is a very important step in social network analysis (Gimenez et al., 2019; Hoppitt & Farine, 2018). To our knowledge, this is the first application of the AME approach in an ecological context. To create and visualize our networks we used the packages Amen (Hoff, 2018), Asnipe (Farine, 2013), gdata (Warnes 2017) and igraph in R (Csardi, Gabor; Nepusz Tamas, 2006).

### Are there social bonds that persist over time?

We investigated if individuals create social bonds that persist over time longer than one breeding occasion by means of two approaches i) contingency tables and ii) the inclusion of time dependent regression terms in the AME modelling framework (Minhas et al., 2016) (see previous section). To answer this question, we analysed data of the period of stability, from 2002 to 2011, dividing this period into two sub-periods of five years (2002-2006 and 2007-2011). In the contingency table approach we tested if the probability of breeding together at least once during the second sub-period was independent of having bred together before at least once in the first period. We built a 3×3 table of frequencies, showing the frequencies of two individuals breeding or not together at least once during the second sub-period depending on if they bred together or not at least once during the first sub-period. We included a third column/line for unknown relationships, i.e. with no information for one or the two of the individuals for that period. We then tested for deviation of random frequencies by Chi Square test.

With the SNA approach we analysed the social bond between individuals using the AME function provided in the Amen package in R and including data from the first period (five previous years) as predictors of association during the second sub-period. We considered that this time window was not too large to include important death events, but large enough to account for the imperfect detection of individuals. To achieve convergence, we increased the number of iterations to 100,000 from the default value of 10,000 and lengthened the burn-in period to 500.

### Had perturbations affected social cohesion in this species?

To assess if perturbations affected social structure in this species, we analysed as previously, with both the contingency table approach and the SNA approach, the social bonding during the period of “transition phase to collapse” (2012-2017). To do so, we tested if the probability of breeding together in this phase (2012-2017) was independent of having bred together during five previous years (2007-2011). We then compared these results from those previously observed during the “stability phase”.

## Results

We analysed a total of 1,610,922 dyadic interactions during the first period (2002-2011) and 368,142 during the second period (2012-2017) (Figure 3). When assessing the social bonding with the contingency table approach, the assumption that breeding aggregations in Audouin’s gull were at random was rejected, and those individuals that bred together during the sub-period 2002-2006 had a higher probability of breeding together during the sub-period 2007-2011 (χ^2^ = 64.685, 1 df, P value < 0.0001).

**Figure 3.**
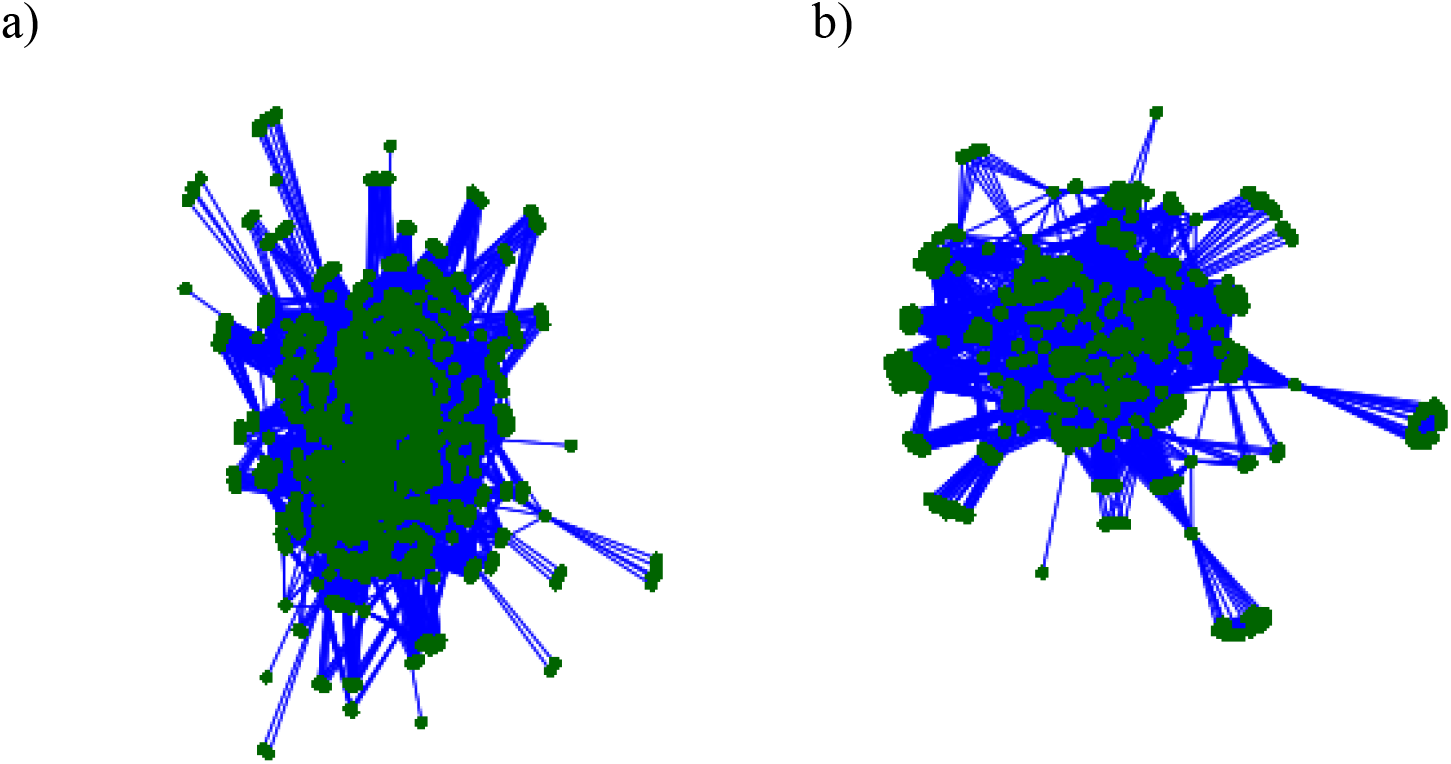
Graphical representation of social networks by the association between individuals of Audouin’s gulls in breeding patches comparing a) the stability phase (2002-2011) and b) the transition phase to colony collapse (2012-2017) (see Figure 1). Each node represents an individual and each edge links those individuals that have bred together in the same patch.

Accordingly, when assessing the social bond with dependent regression terms in the AME function, we show that the probability of breeding together during the second sub-period (2007-2011) depended on whether they have bred previously together in the first sub-period (2002-2006), with a statistically significant coefficient of regression parameter (Table 1; Figure 3).

When we analysed the social bonding during the transition to collapse phase, we observed that the probability of breeding together during the period 2012-2017 did not depend on if they have bred together the five previous years (χ^2^ =1.957, p-value = 0.162) and we could not reject the hypothesis of a random association between individuals. Also the SNA approach showed that breeding aggregations in Audouin’s gull during the transition phase did not depend on if they have bred together the five previous years (Table 1; Figure 3).

**Table 1.** Results of the AME regression function to test if there were social bonds between individuals while breeding during the stable period and during the transition phase to collapse period. The alternative hypothesis is that individuals aggregate annually at random for breeding. “.dyad”: coefficient of the dependent regression term considering the previous dyadic relationship between individuals; “pmean”: posterior mean estimate; “psd”: posterior standard deviation; “z-stat “: nominal z-score.

|  | Stable period |  |  |  | Transition to collapse period |  |  |  |
| --- | --- | --- | --- | --- | --- | --- | --- | --- |
|  | pmean | psd | z-stat | p-value | pmean | psd | z-stat | p-value |
| **intercept** | -1.467 | 0.087 | -16.808 | 0.000 | -0.231 | 0.018 | -12.629 | 0.000 |
| **.dyad** | 6.420 | 0.328 | 19.576 | 0.000 | -0.004 | 0.022 | -0.188 | 0.851 |

## Discussion

By studying a particular ecological system of a colonial long-lived species that experiences a perturbation regime, we showed that social bonding among individuals persist over the years and that perturbations may decrease social cohesion in animal populations.

The characteristic breeding behaviour of the study species that aggregates in patches that change annually, allowed us to show that individuals do not annually breed aggregated at random but rather there is some group stability, with individuals establishing social bonds that persist over time. Group stability can emerge as a product of network self-organization, but may provide the necessary conditions for the evolution of other social processes (Cantor & Farine, 2018; Cantor et al., n.d.). Our results would support the idea that social aggregation during breeding would provide other advantages than the mere defence against predators (Anderson & Hodum, 1993; Oro, 1996), such as social information sharing. Social information sharing is crucial for decision-making in risky behaviours, such as dispersal, and previous studies showed that the perturbed regime in this site caused dispersal to other sites, including colonization of new habitats (Payo-Payo et al., 2017, 2018). Our results suggest that sociality has played a major role driving dispersal and thus populations dynamics in this species, both during the exponential growth after colonization and the collapse after the perturbation regime.

The importance of social information compared to private information is larger under perturbations, even when the quality of social information does not increase compared to a non-perturbed regime (Arganda, Pérez-Escudero, & Polavieja, 2012; Pérez-Escudero & Polavieja, 2017). Under stress conditions, sociality may operate through feedback loops such as social copying for dispersal, causing non-linear population dynamics and playing a critical role on the resilience of populations. We showed here that after a perturbation, not only the number of individuals in the population may decrease (by increased mortality or dispersal) but also its social cohesion, reducing the amount of social information available for those individuals that remain. The perturbation regime suffered by this population has likely triggered a social transition (Pruitt et al., 2018) in collective behaviour from philopatric to dispersal and with the fast diffusion of innovations such as the colonization of harbours, a habitat safe from predators never occupied before (Payo-Payo et al., 2017). Previous studies have shown that responses of populations to perturbations may also depend on individual personalities in the population (Dall, Houston, & McNamara, 2004; Doering, Scharf, Moeller, & Pruitt, 2018; Wolf, van Doorn, Leimar, & Weissing, 2007). For example individual personalities have recently been shown to influence dispersal (Clobert, Le Galliard, Cote, Meylan, & Massot, 2009; Cote, Clobert, Brodin, Fogarty, & Sih, 2010; Fogarty, Cote, & Sih, 2011). Heterogeneities in personalities for dispersal decision-making may have also played a role in this population, with most individuals dispersing to other sites after a period of disturbance, while some individuals remaining philopatric. This probably would have led to a change in the individual personalities composition of the population. This change may have also further consequences, as performance in social systems may improve with heterogeneity in individual personalities in the social group (Fogarty et al., 2011; O’Shea-Wheller, Masuda, Sendova-Franks, & Franks, 2017).

This study opens new research questions about resilience of populations under perturbations; if perturbations are able to affect social cohesion and heterogeneity in personalities in the population, would the perturbed populations be equally resilient to future perturbations? Additionally, in our study population, sociality seemed to operate not right after the first perturbation episode but after a period of maintained perturbations (Payo-Payo et al., 2017); would the type of perturbation, either pulse or in regime (Nimmo, Mac Nally, Cunningham, Haslem, & Bennett, 2015) influence the response of social groups?

We have shown here that perturbations may decrease social cohesion in animal populations, but further studies should be carried out to improve our understanding on the demographic consequences of the breaking down of social bonds under perturbations for population dynamics and resilience on social species.

## Acknowledgements

We would like to thank all the people who have helped with the fieldwork in the Ebro delta over the years, particularly Albert Bertolero, Julia Piccardo, Toni Curcó and the technical staff and volunteers at the Ebro Delta Natural Park. We would also like to thank Peter Hoff, for solving an analytical problem that we encountered at the beginning of the analyzes with the AMEN R package. We also thank the Regional Government of Catalonia, for permits to access the study sites. Elisabeth Rochon corrected the English. Funding came from the Spanish Ministry of Science (CGL2017-85210), Consell Insular de Menorca and grant PICS INTERACT n°07699 (2016, CSIC-CNRS). MG was partially supported by the European Union (MINOUW Project, H2020-634495).

## Authors contributions

MG conceived the idea; MG, OG, RP and RC designed methodology; MG, DO and AB collected the data; MG analysed the data; MG led the writing of the manuscript. All authors critically contributed to the drafts and gave final approval for publication.

## Data accessibility

Data is available via CSIC repository.

