## Supporting information for "Changes in social cohesion in a long-lived species under a perturbation regime"

**Figure S1.** Example of annual patch aggregations at Punta de la Banya colony during the first year of the study period (2002) and de last year (2017).

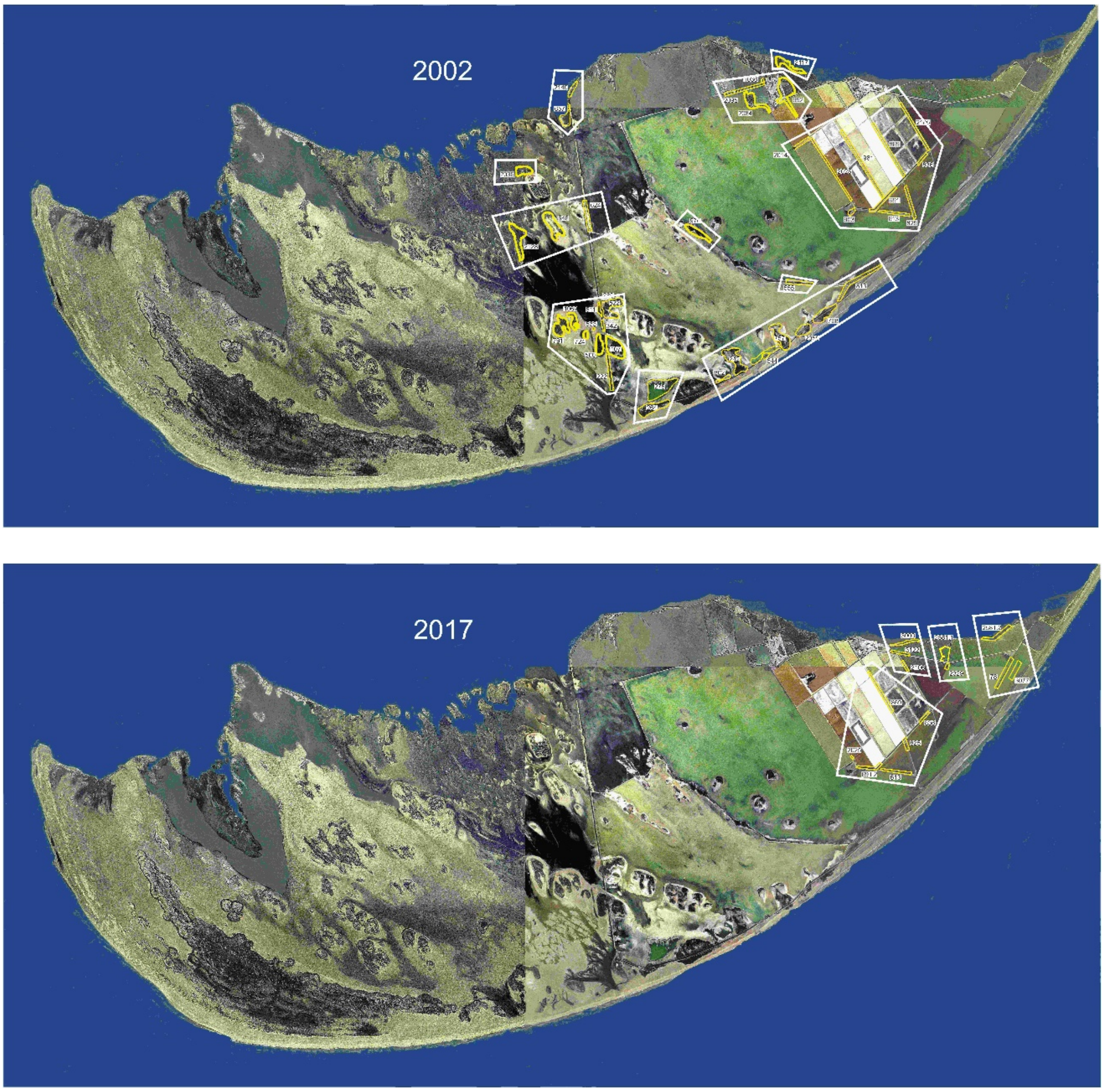

**Figure S2**. Annual number of breeding pairs at the three colonies of the Ebro Delta since colonization of the area in 1981.

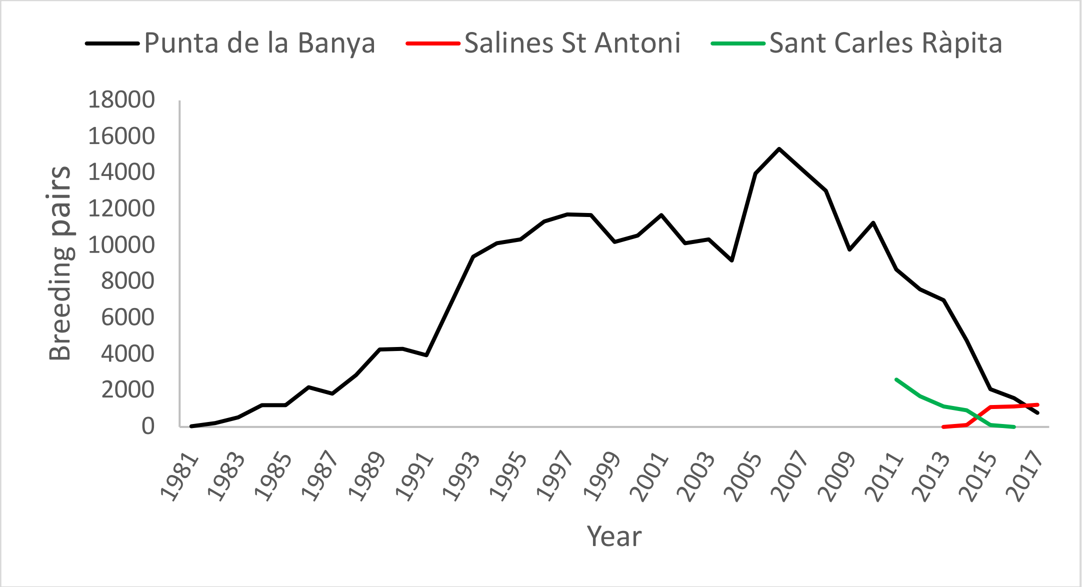
